# Apelin Stimulation of the Perivascular MuSC Niche Enhances Endogenous Repair in Muscular Dystrophy

**DOI:** 10.1101/2022.09.09.507274

**Authors:** Emmeran Le Moal, Yuguo Liu, Jasmin Collerette-Tremblay, Simon Dumontier, Joël Boutin, Junio Dort, Zakaria Orfi, Joris Michaud, Hugo Giguère, Alexandre Desroches, Kien Trân, François Vézina, Sonia Bedard, Catherine Raynaud, Frederic Balg, Philippe Sarret, Michelle S. Scott, Jerome N. Feige, Jean-Bernard Denault, Nicolas A. Dumont, Eric Marsault, Mannix Auger-Messier, C. Florian Bentzinger

## Abstract

Impaired skeletal muscle stem cell (MuSC) function has long been suspected to contribute to the pathogenesis of muscular dystrophy (MD). Here we describe that defects in the endothelial cell (EC) compartment of the perivascular stem cell niche in three different types of MD are associated with inefficient mobilization of MuSCs following tissue damage. Using chemoinformatic analysis, we identified the 13 amino acid form of the peptidic hormone apelin (AP-13) as a candidate for systemic stimulation of skeletal muscle ECs. In dystrophic mice, administration of AP-13 generates a pro-myogenic EC-rich niche that supports MuSC function and markedly improves tissue regeneration, muscle strength, and physical performance. Moreover, we demonstrate that EC specific knockout of the AP-13 receptor leads to regenerative defects that phenocopy major pathological features of MD. Altogether, we provide *in vivo* proof-of-concept that enhancing endogenous repair by targeting the perivascular niche is a viable therapeutic avenue for MD and characterize AP-13 as a novel drug candidate for systemic treatment of stem cell dysfunction.

## Main

Muscular dystrophies (MDs) are a heterogeneous group of rare genetic diseases usually diagnosed in children that are often characterized by severe skeletal muscle wasting and a progressive loss of ambulation. The heterogeneity of genetic defects causing MD represents a major challenge for the development and manufacturing of genome targeted therapeutics. Therefore, mutation-independent treatment approaches that are applicable to a broad spectrum of MD patients represent an important therapeutic opportunity.

Several non-genetic approaches for the treatment of MD have been described. Most common are glucocorticoids, which preserve muscle strength and prolong ambulation in kids with Duchenne MD by reducing inflammation and fibrosis^1^. In addition, a number of drugs that modulate autophagy, reduce oxidative stress, and boost mitochondrial function have been shown to improve myofiber survival and thereby slow disease progression^2^. In preclinical models of certain forms of congenital MD (CMD), anti-apoptotic agents display beneficial effects and are now evaluated in clinical trials^3,4^. Next to targeting inflammation and increasing myofiber survival, stimulating endogenous repair of skeletal muscle has emerged as a therapeutic approach for MD^5^. Myofibers in dystrophic muscles undergo constant cycles of de- and regeneration. Over time, the muscle stem cell (MuSC) pool and its myogenic support cell environment fail to keep up with tissue repair leading to a progressive loss of contractile function^6-8^. Thus, stimulating MuSCs so that myogenic repair outpaces tissue degeneration represents an attractive strategy to preserve functional muscle mass in MD.

In the mdx mouse model of Duchenne MD, increasing asymmetric MuSC divisions using epidermal growth factor promotes the production of committed progeny for differentiation^9^. This strategy leads to enhanced endogenous repair and increases force production in dystrophic muscles. Supplying antibodies or small molecular compounds that activate integrin signaling represents another approach that has been shown to ameliorate the phenotype of mdx mice^10,11^. Increased integrin activity improves the regenerative function of MuSCs and, at the same time, decreases myofiber membrane damage leading to stronger muscles. Altogether, these studies demonstrate that in the mdx mouse model, multiple approaches that directly target MuSC function have therapeutic effects. Maintenance, self-renewal, and differentiation of MuSCs are tightly controlled by their ECM environment and different supportive cell populations in the stem cell niche^12^. Our recent work has shown that a reduction of inflammatory cells in the MuSC niche using pro-resolving mediators mobilizes the stem cell pool and increases myogenic differentiation promoting myofiber repair in mdx mice^13^. These results suggest that strategies aiming at a normalization of the stem cell microenvironment represent an additional intervention point for stimulating endogenous repair in Duchenne MD.

In contrast to Duchenne MD that is caused by mutations in the inner membrane protein dystrophin, most forms of CMD are caused by truncation or absence of extracellular matrix (ECM) proteins or their receptors^14^. Interestingly, regenerative defects have been described in mouse models of both laminin-a2 (LAMA2) MD and collagen VI (ColVI)-related myopathy^15,16^. Loss of ECM proteins in certain forms of CMD leads to the upregulation of partially compensating alternative isoforms^17,18^. In addition, severe forms of CMD lead to major shifts in the cellular composition of the tissue, including increased amounts of inflammatory and pro-fibrotic cells^19,20^. Therefore, changes in the cellular and extracellular niche environment of MuSCs likely have a dominant role in triggering regenerative defects in CMDs. These observations support the notion that stimulating MuSCs or targeting the niche to normalize stem cell function and enhance endogenous repair also represent a potential therapeutic avenue to slow disease progression in this group of MDs.

Here we systematically investigated MuSC function and pathologic adaptations of the stem cell niche in mouse models of Duchenne MD, LAMA2 MD, and ColVI-related myopathy. We demonstrate that an impaired expansion capacity of the stem cell pool and aberrant changes in the endothelial cell (EC) compartment of the niche are a common denominator of all three types of MD. Using chemoinformatic screening, we identify the 13 amino acid hormone apelin as a systemically administrable therapeutic that stimulates ECs and enhances MuSC function in dystrophic muscles. Lastly, we demonstrate that EC specific loss of the apelin receptor (APJ) leads to aberrant changes in the skeletal muscle microvasculature and phenocopies major features of MD. In summary, our work characterizes defects in the perivascular MuSC niche as a hallmark of MD and identifies apelinergic signaling as a therapeutic target for the stimulation of endogenous repair.

## Results

### Muscular dystrophy affects the proliferative capacity of MuSCs

To systematically assess stem cell function and the regenerative capacity of skeletal muscle across a spectrum of different types of MD, we compared mouse models of ColVI-related myopathy (d16), Duchenne MD (mdx), and LAMA2 MD (dyW), to wild-type (wt) animals (**Fig. 1a**). In order to maximize tissue regeneration and MuSC activation, we injected mice with the snake venom cardiotoxin (CTX) and analyzed the tissue at 5 and 10 days post injury (dpi). Hematoxylin and eosin staining of muscle cross-sections revealed that at 5 and 10 dpi all three MD models displayed changes in tissue architecture including an increased interstitial volume and a higher abundance of mononuclear cells when compared to wt controls (**Fig. 1b**). In the uninjured baseline, differences in fiber size were observed in all dystrophic models compared to control animals (**Fig. 1c**). mdx and d16 did not display differences in fiber size at 10 dpi (**Fig. 1d**). In contrast, the size distribution at 10 dpi was shifted significantly towards smaller fibers in dyW mice. Immunostaining for embryonic myosin heavy chain (eMHC), a marker of newly formed muscle fibers, showed that, following injury, tissue maturation in d16 and dyW mice was delayed compared to wt controls and remained increased with respect to the baseline in uninjured muscles (**Fig. 1e-i**). Moreover, d16 and dyW mice displayed an increased abundance of the fibrosis marker fibronectin at both 5 and 10 dpi (**Fig. 1e, 1j-m**). Staining for the MuSC marker Pax7 identified reduced stem cell numbers in all three MD models at 5 dpi, and in d16 and dyW mice at 10dpi (**Fig. 1n-q**). Importantly, relative to pre-injury levels, MuSC numbers in the dystrophic mouse models did not increase by the same magnitude as in wt mice (**Fig. 1r**). The MuSC pool in dyW mice showed a particularly impaired expansion potential and did not change significantly compared to pre-injury levels. Thus, a reduced capacity for mobilization of the stem cell pool is a feature of multiple types of MD. Overall, the severity of the regenerative phenotype increases in from mdx to d16 mice and is most pronounced in the dyW model (**Fig. 1s**).

**Fig. 1:**
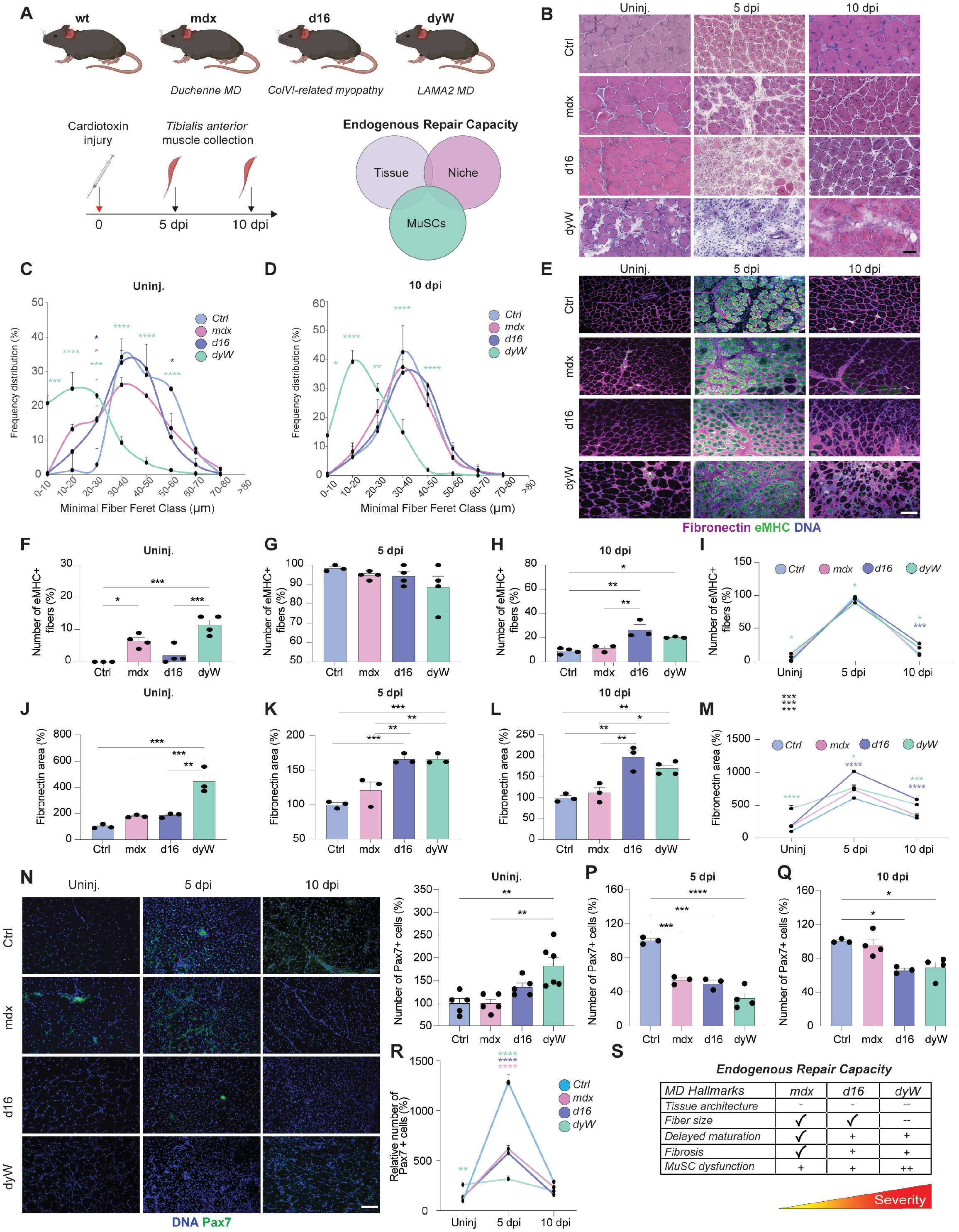
Muscular dystrophy (MD) affects the proliferative capacity of MuSCs. **a**, Scheme outlining the MD models used in our study and graphical overview of the experimental timeline. d16 mice are a model for ColVI-related myopathy, mdx mice for Duchenne MD, and dyW mice for LAMA2 MD. C57BL/6 mice were used as background matched wild-type (wt) controls. **b**, Representative haematoxylin and eosin stained cross sections of the *tibialis anterior* (TA) muscle from wt controls (ctrl), mdx, d16 and dyW mice under uninjured (uninj.) and at 5 and 10 days post cardiotoxin injury (dpi). **c**,**d**, Frequency distribution of minimal fiber feret classes in TA muscles of ctrl, mdx, d16 and dyW muscles under uninjured conditions (c) and at 10 dpi (d). **e-m**, Immunostaining and quantification of embryonic myosin heavy chain positive fibers (eMHC) co-stained with fibronectin in TA muscle sections of ctrl, mdx, d16 and dyW mice under uninjured conditions (f,j), and at 5 (g,k) and 10 (h,l) dpi. Kinetics of eMHC positive fibers (i) and fibronectin expression over the regenerative time course (m). **n-r**, Immunostaining and quantification of Pax7 positive cells in TA muscle sections in ctrl, mdx, d16 and dyW mice under uninjured conditions (o), and at 5 (p) and 10 dpi (q). Kinetics of Pax7 positive cells over the regenerative time course (r). **s**, Table summarizing the severity of endogenous repair defects in dystrophic mouse models. - = decreased, - - strongly decreased, + = increased, + + strongly increased, √ = similar to ctrl. Results are expressed as means + sem. n ≥ 3 mice per condition. Scale bars = 50 μm (b) and 100 μm (e and n). P values were calculated using one and two-way ANOVA with Tukey’s post-hoc test. \**P*<0.05, \*\**P*<0.01, \*\*\**P*<0.001, \*\*\*\**P*<0.0001.

### MD affects microvascular remodeling

In order to study the cellular composition of the MuSC niche in the different types of MD, we quantified the number of fibro–adipogenic progenitors (FAPs), macrophages, and ECs under uninjured conditions, and at 5 and 10 dpi. Staining for Pdgrfα revealed increased FAP numbers at baseline in dyW mice (**Fig. 2a,b**). At 5 dpi, FAP numbers increased in mdx and dyW mice, while they decreased in d16 muscles compared to wt controls (**Fig. 2c,e**). In dyW mice, FAP numbers remained higher than in wt controls at 10 dpi (**Fig. 2d**). F4/80 positive macrophages showed an increased abundance at baseline and following injury in all MD models (**Fig. 2f-j**). Compared to wt mice, mdx mice showed a particularly pronounced macrophage response at 5 dpi. Notably, respective to wt control muscles, the number of ECs was lower in mdx and dyW mice at baseline (**Fig. 2k,l**). Moreover, EC numbers were dramatically reduced at both 5 and 10 dpi in all three MD models (**Fig. 2k,m-o**). These results support the notion that impaired microvascular remodeling is a pathologic hallmark of MD.

**Fig. 2:**
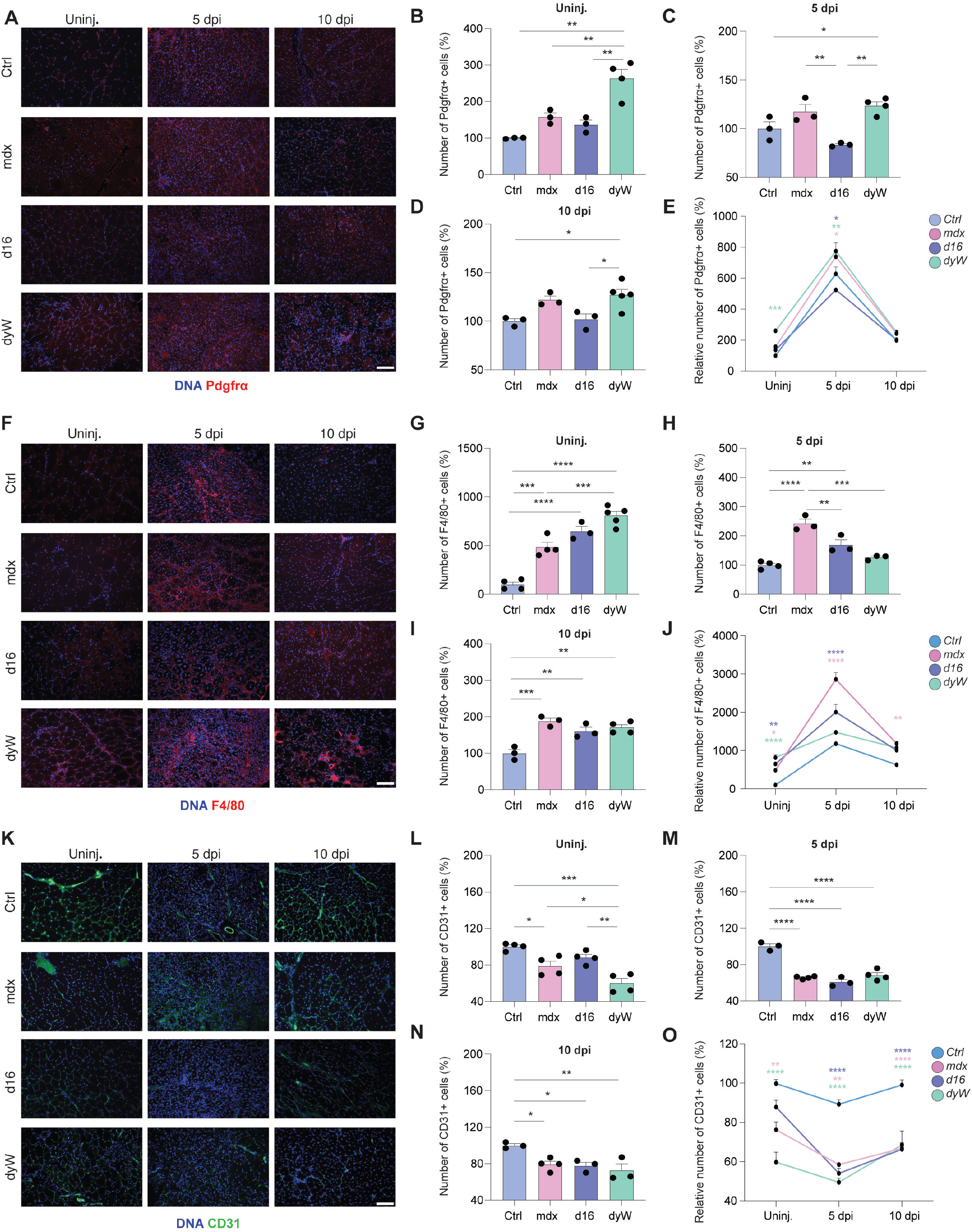
MD affects microvascular remodeling. **a-e**, Immunostaining and quantification of Pdgfrα positive cells in TA muscle sections in ctrl, mdx, d16 and dyW mice under uninjured conditions (b), and at 5 (c) and 10 dpi (d). Kinetics of Pdgfrα positive cells over the regenerative time course (e). **f-j**, Immunostaining and quantification of F4/80 positive cells in TA muscle sections in ctrl, mdx, d16 and dyW mice under uninjured conditions (g), and at 5 (h) and 10 dpi (i). Kinetics of F4/80 positive cells over the regenerative time course (j). **k-o**, Immunostaining and quantification of CD31 positive cells in TA muscle sections in ctrl, mdx, d16 and dyW mice under uninjured conditions (l), and at 5 (m) and 10 dpi (n). Kinetics of CD31positive cells over the regenerative time course (o). Results are expressed as means + sem. n ≥ 3 mice per condition. Scale bars 100 μm. P values were calculated using one and two-way ANOVA with Tukey’s post-hoc test. \**P*<0.05, \*\**P*<0.01, \*\*\**P*<0.001, \*\*\*\**P*<0.0001.

### Identification of AP-13 as a skeletal muscle EC stimulatory molecule

In light of microvascular phenotype we observed in dystrophic muscles, we set out to identify angiogenic factors with the potential to stimulate ECs. G protein-coupled receptors (GPCRs) represent the largest family of druggable targets in the human genome^21^. Based on the single cell atlas by De Micheli et al.^22^, we compiled a list of GPCRs expressed by skeletal muscle ECs under uninjured conditions and at 5 dpi (**Fig. 3a,b**). We observed that APJ, the receptor for the small peptidic hormone apelin, shows the second highest expression at 5 dpi. Mapping of APJ to the whole single cell transcriptome of skeletal muscle showed that its expression is highly specific to ECs and does not overlap with Pax7 positive MuSCs (**Fig. 3c**). APJ immunostaining of skeletal muscle sections of wt mice at 5 dpi confirmed a distinct colocalization with ECs, while MuSCs, macrophages, and FAPs did not express discernable levels of the receptor (**Fig. 3d,e**).

The APJ ligand apelin is naturally produced as a 77-amino-acid precursor that is processed into active 36, 17, and 13 amino acid fragments found in the systemic circulation^23^. Apelin 13, the smallest active form of apelin, has a molecular weight of 1.5 kDa and is naturally pyroglutamylated at its N-terminus (**Fig. 3f,g**). To investigate potential stimulatory effects on ECs, we produced the pyroglutamylated apelin 13 (AP-13) fragment using solid phase peptide synthesis (**Supplementary Fig. 1a**)^24^. Purity of AP-13 was confirmed to be >95% using analytical UPLC/MS (data not shown). *In vitro* experiments revealed that AP-13 elicits a dose dependent proliferative response of ECs (**Fig. 3h**). In contrast, MuSC derived primary myoblasts did not react with increased proliferation to AP-13 (**Fig. 3i**). Altogether, these results identify AP-13 is a candidate for the targeted stimulation of ECs in dystrophic skeletal muscle.

**Fig. 3:**
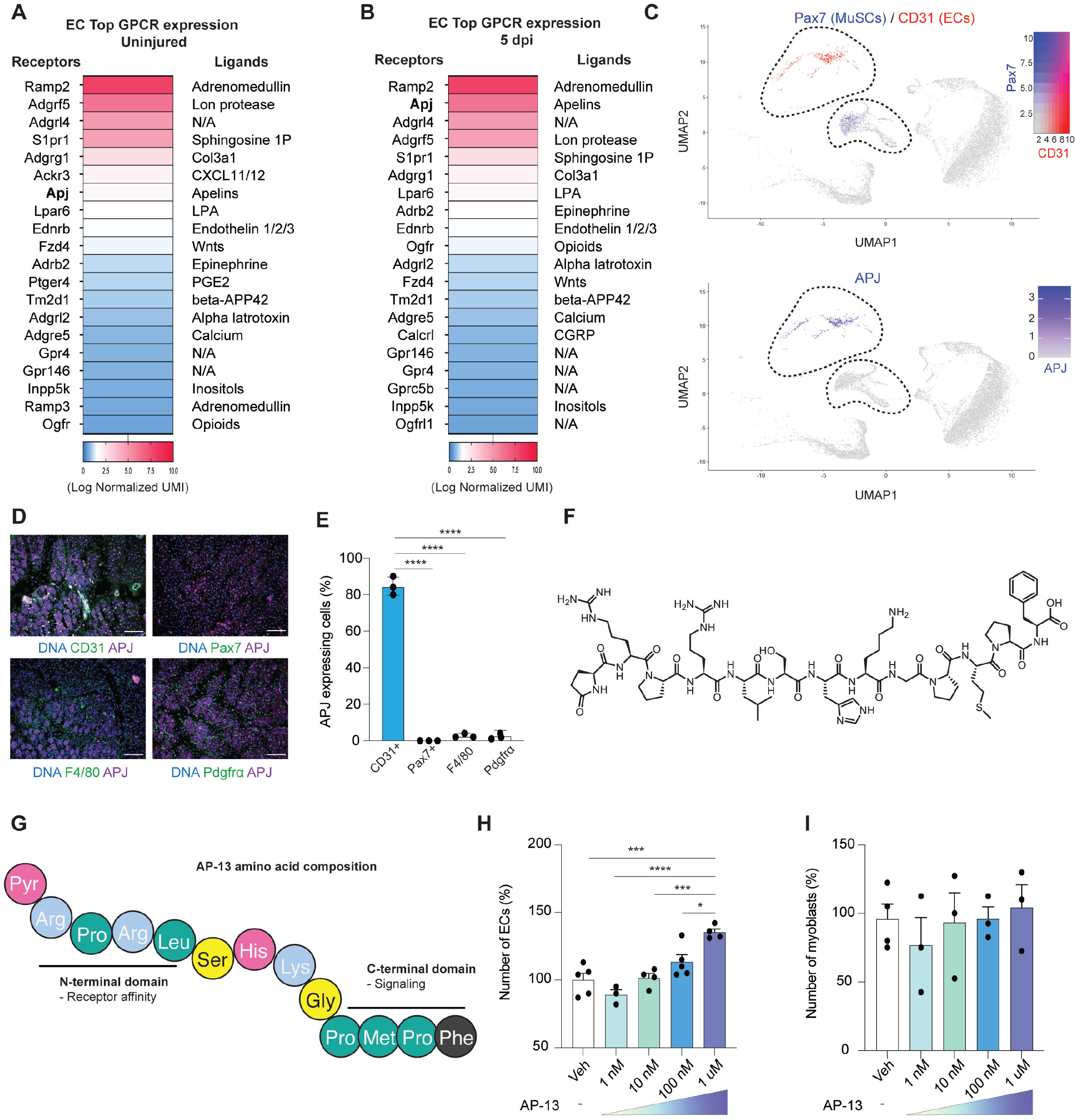
Identification of AP-13 as an EC stimulatory molecule. **a,b**, Heat map of the top 20 most expressed GPCRs in ECs under uninjured conditions (a) and at 5 dpi (b) and their cognate ligands based on single cell transcriptomics of TA muscles. UMI = Unique molecular identifiers **c**, Uniform manifold approximation and projection (UMAP) representation of single cell transcriptomes obtained from TA muscles. Expression of the apelin receptor APJ is shown next to Pax7 to label MuSCs and CD31 to label endothelial cells (ECs). **d,e**, Immunostaining and quantification of APJ levels co-stained with CD31, Pax7, F4/80, and PDGFRα in TA muscle cross sections of wt mice at 5 dpi. **f,g**, Chemical structure (f) and amino acid profile of the pyr-apelin-13 peptide (AP-13) (g). **h,i**, Proliferation of ECs (h) and MuSC derived myoblasts (i) in response to increasing concentrations of AP-13. Results are expressed as means + sem. n ≥ 3 mice per condition. Scale bars = 100 μm. P values were calculated using one way ANOVA with Tukey’s post-hoc test (h,i). \**P*<0.05, \*\**P*<0.01, \*\*\**P*<0.001, \*\*\*\**P*<0.0001.

### AP-13 mediated stimulation of the perivascular niche improves endogenous repair

Since they display the most pronounced regenerative defects and microvascular phenotype, we decided to assess the ability of AP-13 to stimulate skeletal muscle ECs in dyW mice. To this end we implanted the dystrophic mice at the age of 14 days with osmotic pumps supplying AP-13 or PBS vehicle (veh) for four weeks (**Fig. 4a**). As predicted by our *in vitro* results, we observed that AP-13 treatment increased the number of ECs in dyW muscles by 59% when compared to the veh condition (**Fig. 4b**). AP-13 caused a 14% reduction in the number of FAPs but did not affect macrophages (**Fig. 4c,d**). No significant effect on fibrosis or fibers with membrane damage that stain for intracellular IgG were observed as a consequence of AP-13 treatment (**Fig. 4e,f**). Moreover, western blot for cleaved caspase 3 revealed that apoptotic processes were not altered in the AP-13 group (**Supplementary Fig. 2a-c**). Interestingly, AP-13 mediated stimulation of ECs was accompanied by a 58% increase in Pax7 positive MuSCs and an 80% increase of differentiating myogenin (MyoG) positive cells (**Fig. 4g,h**). In agreement with increased MuSC proliferation and differentiation, AP-13 treatment led to a 72% increase in newly formed eMHC positive fibers (**Fig. 4i**). Thus, AP-13 stimulation of skeletal muscle ECs promotes MuSC function and endogenous repair in dyW mice but does not affect fibrosis and the survival or integrity of muscle fibers.

**Fig. 4:**
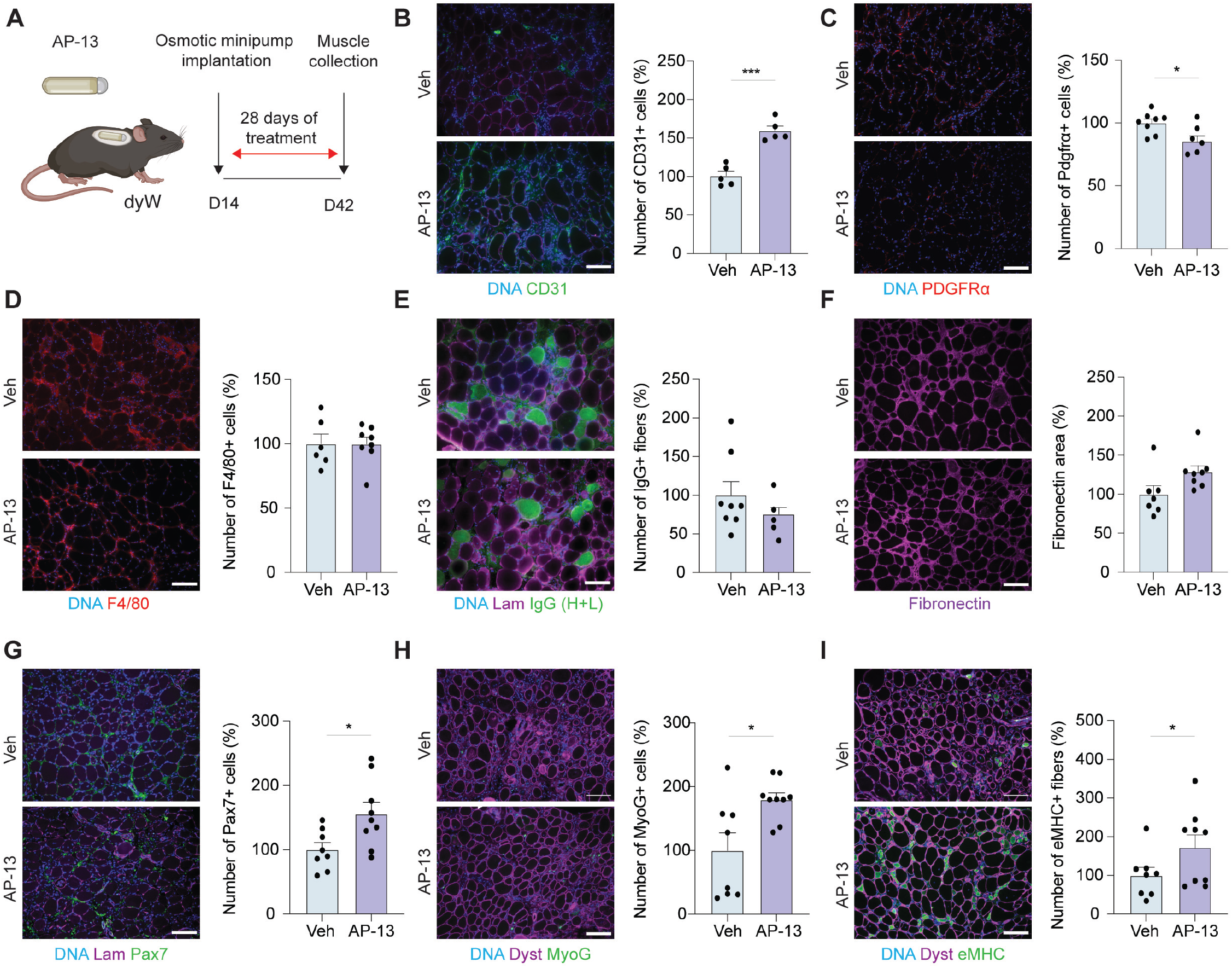
AP-13 stimulates the perivascular MuSC niche and improves endogenous repair in MD. **a**, Scheme outlining the AP-13 treatment strategy of dyW mice. **b**, Immunostaining and quantification of CD31 positive cells in TA muscle cross sections of dyW mice treated with vehicle (veh) or AP-13. **c**, Immunostaining and quantification of Pdgfrα positive cells in TA muscle cross sections of dyW mice treated with veh or AP-13. **d**, Immunostaining and quantification of F4/80 positive cells in TA muscle cross sections of dyW mice treated with veh or AP-13. **e**, Immunostaining and quantification of IgG positive fibers in TA muscle cross sections of dyW mice treated with veh or AP-13. **f**, Immunostaining and quantification of the fibronectin positive area in TA muscle cross sections of dyW mice treated with veh or AP-13. **g**, Immunostaining and quantification of Pax7 positive cells in TA muscle cross sections of dyW mice treated with veh or AP-13. **h**, Immunostaining and quantification of Myogenin (MyoG) positive cells in TA muscle cross sections of dyW mice treated with veh or AP-13. **i**, Immunostaining and quantification of eMHC positive fibers in TA muscle cross sections of dyW mice treated with veh or AP-13. Results are expressed as means + sem. n ≥ 3 mice per condition. Scale bars = 100 μm. P values were calculated using students *t*-test. \**P*<0.05, \*\**P*<0.01, \*\*\**P*<0.001, \*\*\*\**P*<0.0001.

### Systemic AP-13 treatment slows disease progression in MD

Cumulation over the four-week treatment course revealed that the AP-13 treated group of dyW mice displayed a 12% higher average body weight (**Fig. 5a,b**). As opposed to veh treated dyW mice, not a single animal in the AP-13 group died before the study endpoint (**Fig. 5c**). To uncouple potential positive effects of AP-13 on skeletal muscle in dyW mice from effects on secondary tissues, we performed *ex-vivo* and *in situ* muscle force measurements (**Fig. 5d,e)**. This revealed that *extensor digitorum longus* (EDL) muscles isolated from dyW mice that were treated for 3 weeks with AP-13 were in average 109% stronger than muscles from the veh control group (**Fig. 5f**). Similarly, *in situ* stimulation of the posterior muscle compartment of the lower leg revealed a 69% increase in force production in the AP-13 group compared to the veh condition (**Fig. 5g**).

**Fig. 5:**
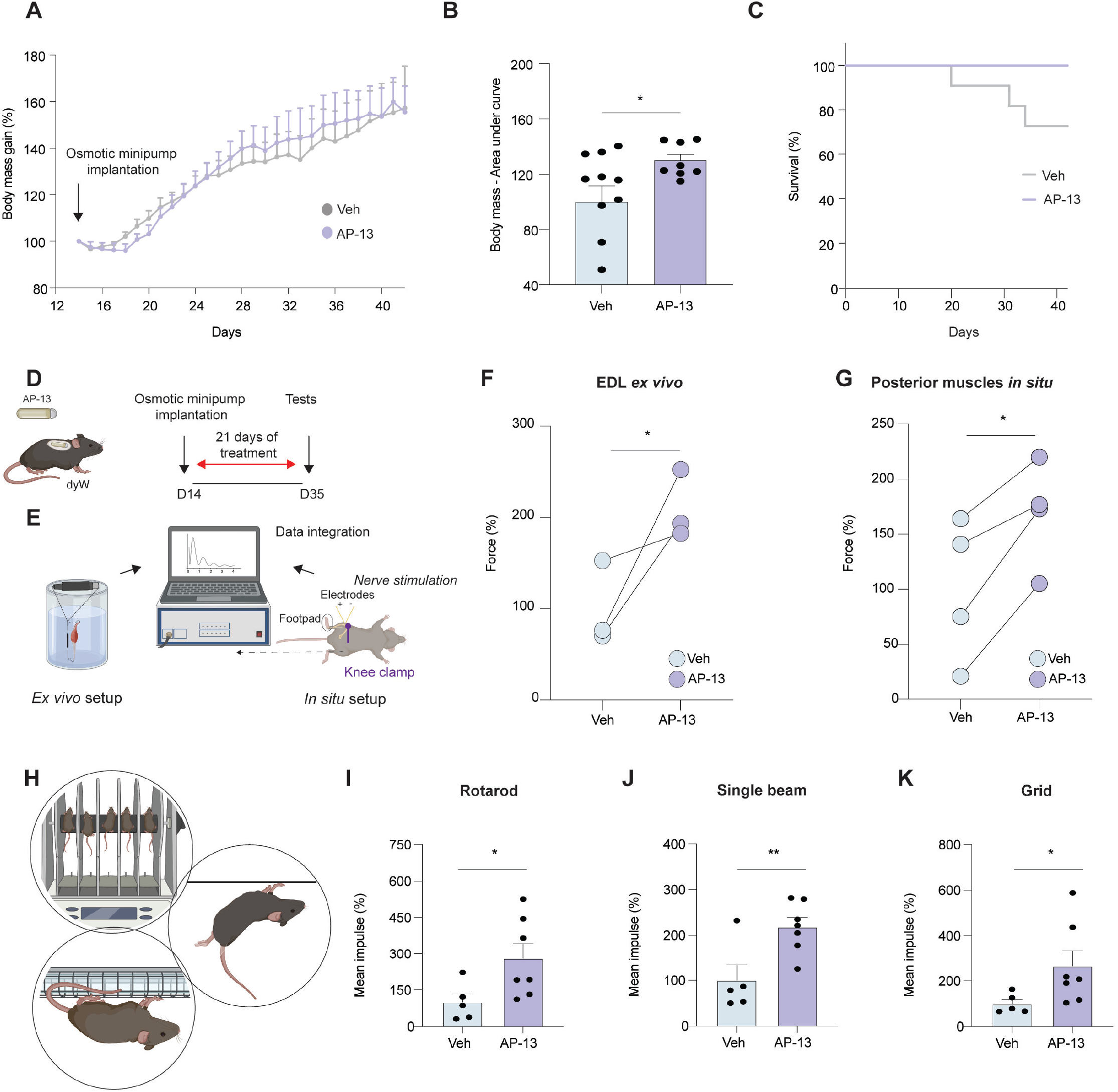
Systemic AP-13 treatment slows disease progression in MD. **a**, Body weight evolution of dyW mice over the veh or AP-13 treatment time-course. **b**, Cumulative body weight gain of veh and AP-13 treated dyW mice. **c**, Survival curve of veh or AP-13 treated dyW mice over the treatment time-course. **d,e**, Scheme outlining the treatment strategy of dyW mice with veh or AP-13 (d) and the set-up for *ex-vivo* and *in situ* muscle force measurements (e). **f**, Quantification of normalized *ex-vivo* force of extensor digitorum longus muscles of dyW mice treated with veh or AP-13. Black lines designate littermates. **g**, Quantification of normalized *in situ* isometric torque force of posterior muscles of the lower leg of dyW mice treated with veh or AP-13. Black lines designate littermates. **h**, Scheme depicting the rotarod, single beam suspension, and grid suspension fitness tests. **i-k**, Quantification of the normalized mean impulse (time to task failure x body mass) of dyW mice treated with veh or AP-13 in the rotarod assay (i), single beam challenge (j) and horizontal grid test (k). Results are expressed as means + sem. n≥8 (a-c), n≥3 (f-g), and n≥5 (i-k) mice per condition. P values were calculated using students *t*-test. \**P*<0.05, \*\**P*<0.01, \*\*\**P*<0.001, \*\*\*\**P*<0.0001.

To determine whether AP-13 treated dyW mice show a global amelioration of disease progression compared to the veh group, we challenged them using a number of physical performance tests. This revealed a 180%, 118% and 166% increase in the mean impulse in AP-13 treated mice in the rotarod, single beam, and horizontal grid challenge respectively when compared to veh treated animals (**Fig. 5h-k**). In summary, AP-13 mediated stimulation of endogenous repair in dyW muscles is accompanied by dramatic gains in muscle force and overall physical performance.

### AP-13 treatment causes no adverse cardiac effects

MDs are frequently accompanied by cardiac dysfunction. Normalized to their body weight, dyW mice in the veh group had 30% heavier hearts than wt mice (**Supplementary Fig. 3a,b**). This suggests the presence of hypertrophic compensatory growth or fibrosis. Compared to veh, AP-13 reversed this phenotype and the treated dyW mice had heart weights that were not different from the untreated wt group. Moreover, echocardiography revealed that the cardiac index and fractional shortening of hearts in the AP-13 treated group were similar to untreated wt hearts, while they were increased by 76% and 20% respectively in the veh group (**Supplementary Fig. 3c,d**). Therefore, we conclude that AP-13 has no adverse effects on heart function.

### EC specific knockout of APJ phenocopies MD features

Our results suggest that AP-13 not only holds therapeutic potential for MD but, given the high expression of APJ in skeletal muscle ECs, is also an important endogenous regulator of angiogenesis in this tissue. Indicative of an angiogenic response induced by the chronic de- and regenerative processes in dystrophic muscle, we observed a 71% upregulation of APJ in the microvasculature of uninjured dyW mice when compared to wt controls (**Supplementary Fig. 4a**). To address the role of endogenous APJ in ECs, we generated mice carrying a CreERT2 cassette under the Cdh5 promoter with floxed alleles of APJ (APJ^EC^KO) and, following tamoxifen mediated gene excision, analyzed them at 5 and 10 dpi (**Fig. 6a,b**). Loss of APJ led to a 37% and 41% reduction in CD31 positive cells at 5 and 10 dpi respectively when compared to the wt condition (**Fig. 6c-e**). Interestingly, hematoxylin and eosin staining of skeletal muscle cross sections revealed that APJ^EC^KO mice displayed an increase in mononuclear cells and interstitial volume at both time-points after injury that resembles the regenerative phenotype observed in severe MD (**Fig. 6f**). Compared to wt controls, fiber size was significantly reduced in APJ^EC^KO mice at 10 dpi (**Fig. 6g,h**). Indicating a delayed regenerative response in APJ^EC^KO mice, staining for eMHC showed a 6% decrease in eMHC positive fibers at 5 dpi and a 306% increase at 10 dpi (**Fig. 6i-k**). Fibronectin staining also revealed an 43% increase of the fibrotic area 5 dpi and an increase of 53% at 10 dpi in APJ^EC^KO mice (**Fig. 6l-n)**. Supporting the notion that, similar to MD, microvascular defects in skeletal muscle affect the stem cell pool in APJ^EC^KO mice, the number of Pax7 positive MuSCs was reduced by 50% at 5 dpi compared to wt controls (**Fig. 6o-q**). Moreover, correlating with delayed tissue regeneration, APJ^EC^KO mice showed a 30% reduction of differentiating MyoG positive MuSCs at 5 dpi and a 52% increase at 10 dpi (**Fig. 6r-t**). We conclude that EC specific loss of APJ leads to defective microvascular remodeling, an impaired expansion capacity of the MuSC pool, and a myopathic phenotype reminiscent of MD.

**Fig. 6:**
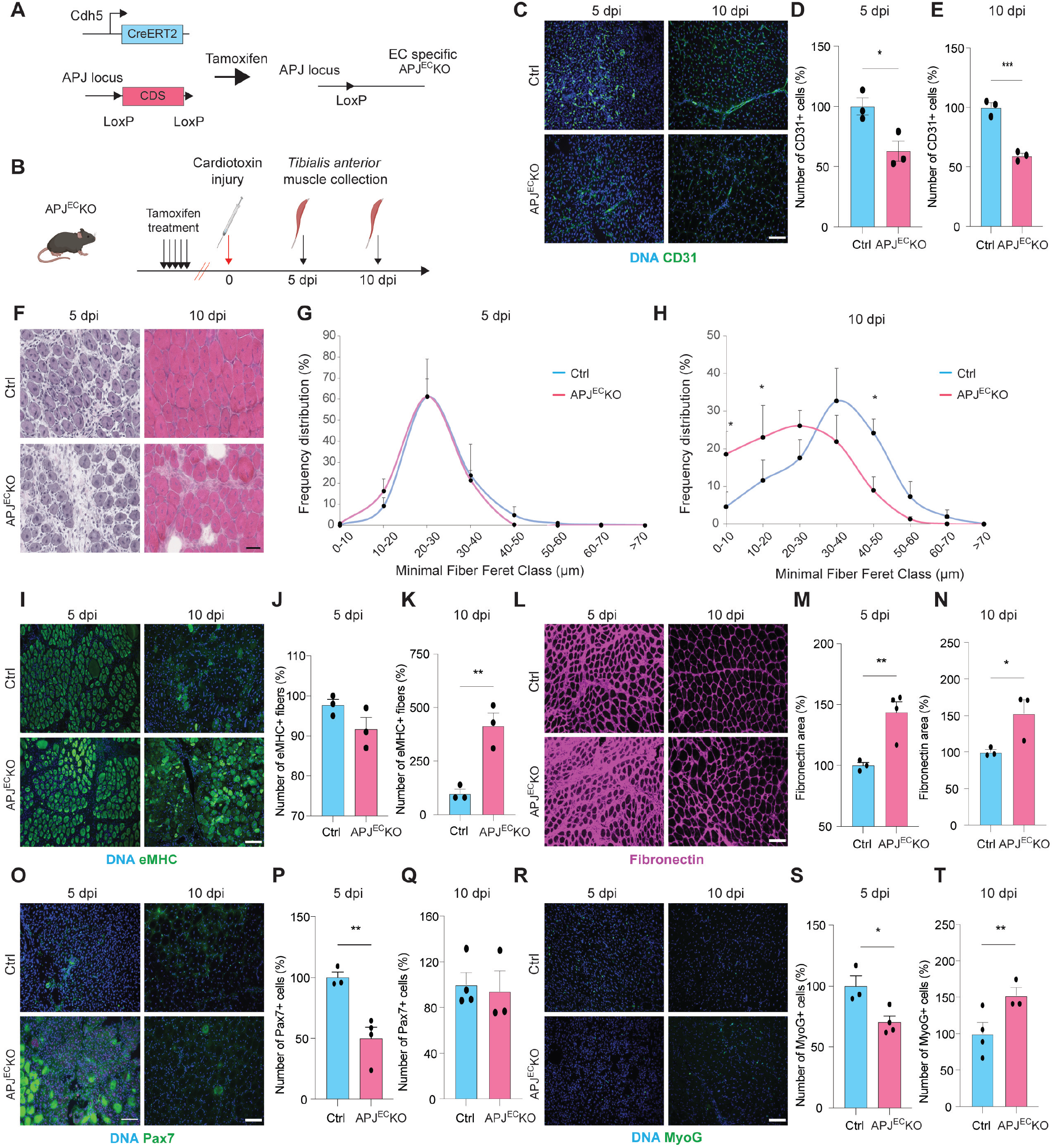
EC specific knockout of APJ phenocopies MD features. **a,b**, Scheme outlining the breeding strategy to generate APJ^EC^KO mice and the muscle injury protocol. **c-e**, Immunostaining and quantification of CD31 positive cells in TA muscle cross sections of wt control (ctrl) and APJ^EC^KO mice at 5 (d) and 10 dpi (e). **f**, Representative haematoxylin and eosin-stained cross sections of the TA muscle of ctrl and APJ^EC^KO mice at 5 and 10 dpi. **g,h**, Frequency distribution of minimal fiber feret classes in TA muscles of ctrl and APJ^EC^KO muscles 5 (g) and 10 dpi (h). **i-k**, Immunostaining and quantification of eMHC positive fibers in TA muscle sections in ctrl and APJ^EC^KO mice at 5 (j) and 10 dpi (k). **l-n**, Immunostaining and quantification of the fibronectin positive area in TA muscle sections in ctrl and APJ^EC^KO mice at 5 (m) and 10 dpi (n). **o-q**, Immunostaining and quantification of Pax7 positive cells in TA muscle sections in ctrl and APJ^EC^KO mice at 5 (p) and 10 dpi (q). **r-t**, Immunostaining and quantification of MyoG positive cells in TA muscle sections in ctrl and APJ^EC^KO mice at 5 (s) and 10 dpi (t). Results are expressed as means + sem. n≥3 per condition. Scale bars = 100μm. P values were calculated using student *t*-test (d,e,j,k,m,n,p,q,s,t) and two-way ANOVA with Tukey’s post-hoc test (g,h). \**P*<0.05, \*\**P*<0.01, \*\*\**P*<0.001, \*\*\*\**P*<0.0001.

Altogether, our study identifies microvascular defects associated with regenerative dysfunction as a major pathologic feature of MD. We identify APJ in ECs as a novel therapeutic target for MD and demonstrate that AP-13 is a viable pharmacologic option for the systemic stimulation of endogenous MuSC mediated repair.

## Discussion

We observed that defective microvascular remodeling is a pathologic hallmark of multiple types of MD that correlates with MuSC dysfunction. Stimulation of angiogenesis using AP-13 is able to enhance MuSC function in dystrophic tissues and leads to a dramatic amelioration of disease progression. Complementing our findings, it has been shown that EC specific knockout of *Flt1* leads to a higher capillary density in skeletal muscle that goes along with improved histological parameters and force generation in mdx mice^25^. The microvasculature supplies tissues with nutrients, circulating growth factors, gases, and electrolytes, and serves as a sink for waste products. Ischemia, the reduction of blood flow to tissues, leads to hypoxia, increased inflammation, adaptations in cellular metabolism that under extreme conditions cause cell death. Ischemia is typically accompanied by an increase in the expression of angiogenic molecules such as vascular endothelial growth factor (VEGF), which promotes the migration of ECs for vascular sprouting^26^. Suggesting an important role of APJ in angiogenesis, loss-of-function models of this receptor show severe EC sprouting defects^27^. Thus, it is conceivable that increased levels of APJ in the vasculature in dystrophic skeletal muscles are part of an ischemic angiogenic response.

Next to their role in providing access to the systemic circulation through the microvasculature, ECs control MuSC function through diverse paracrine factors including IGF1 and angiopoietin 1^12^. Vice-versa, MuSCs stimulate ECs through the release of VEGF^28^. Using elegant tissue clearing methods, Verma et al., have shown that ECs present Notch ligands promoting MuSC self-renewal in the proximity of blood vessels^29^. These data suggest that stimulation of ECs by exogenously supplied AP-13 leads to a feed-forward loop in dystrophic skeletal muscle that amplifies the supportive interplay of these two cell types and thereby boosts the regenerative response.

The most abundant isoforms of apelin found in human plasma are apelin-13 and -17^30^. Plasma concentrations of AP-13 decrease during aging and constitutive global apelin knockout mice display a sarcopenic phenotype^31^. Systemic supplementation of AP-13 reverses diverse hallmarks of skeletal muscle aging including regenerative dysfunction. Interestingly, it has been described that the skeletal muscle microvasculature in sedentary older adults is significantly reduced^32^. These observations suggest that stimulation of the perivascular niche could also contribute to the beneficial effects of AP-13 on endogenous repair and MuSC function in aging. Moreover, physical exercise increases AP-13 plasma concentrations in humans^31^. Thus, the reduced physical activity of dystrophic mice may lead to decreased endogenous levels of AP-13 providing a therapeutic window for supplementation.

Several forms of MD are accompanied by cardiac complications^33^. Our results demonstrate that AP-13 does not negatively affect heart function. Moreover, we observed no changes in histology in diverse organs of AP-13 treated mice (data not shown). Clinical trials that addressed the effects of AP-13 on insulin sensitivity in humans have shown no adverse cardiovascular events and safety reports did not show any other side effects related to the treatment^34^. Thus, AP-13 possesses a promising safety profile as a therapeutic molecule. The half-life of AP-13 *in vivo* is <5 min^35,36^. Importantly, our previous work has shown that macrocyclization and other modifications of apelin 13 can prolong its half-life in plasma *ex-vivo* by more than an order of magnitude and, at the same time, increase its binding affinity for the APJ receptor^37^. Moreover, we demonstrated that substitution of the C-terminal Phe(13) of AP-13 with unnatural amino acids leads to a higher affinity to the APJ receptor and inhibits the production of intracellular cAMP more potently^24^. Thus, we conclude that AP-13 represents an ideal candidate for hit-to-lead optimization of a pharmaceutical compound for the treatment of MD.

Taken together, our study demonstrates that changes in the perivascular niche are accompanied by MuSC dysfunction in MD. We identify and characterize the systemically administrable AP-13 as a therapeutic molecule that is able to increase the EC content in dystrophic muscles and thereby mobilizes endogenous MuSC mediated myofiber repair.

## Acknowledgements

We thank Kristy Red-Horse (Stanford University) for providing the APJ flox mice. C.F.B. is supported by the Canadian Institutes of Health Research (CIHR, PJT-162442), the Natural Sciences and Engineering Research Council of Canada (NSERC, RGPIN-2017-05490), the Fonds de Recherche du Québec - Santé (FRQS, Dossiers 296357, 34813, and 36789), the ThéCell Network (supported by the FRQS), the Canadian Stem Cell Network, and a research chair of the Centre de Recherche Médicale de l’Université de Sherbrooke (CRMUS). E.M. was supported by CIHR (PJT-376770 and PJT-399567), NSERC (RGPIN-1140468), and the Canada foundation for innovation (CFI, 32000). M.A.M. is supported by CIHR (PJT-376770 and PJT-399567), the FRQS (Dossiers 29255 and 284164), CFI (34568), and the Heart and Stroke Foundation of Canada new investigator awards. J.B. is supported by NSERC (RGPIN-2017-05988). P.S. holds the Canada Research Chair in Neurophysiopharmacology of Chronic Pain and is supported by CIHR (FDN-148413). N.A.D is supported by FRQS (Dossiers 35015 and 296512), CIHR (PJT-156408 and PJT-155369), and NSERC (RGPIN-2018-05979). E.L.M. is supported by a postdoctoral fellowship of the FRQS (Dossier 258477). J.C.T. is supported by a Canada Graduate Scholarships-Master’s (CGS M) fellowship and the FRQS (Dossier 254142). H.G. is supported by a CIHR Frederick Banting and Charles Best graduate scholarship (CGS-D).

## Author contributions

C.F.B initiated and managed the project. E.L.M, J.C.T, Y.L., J.D., Z.O., J.M., J.B.D., A.D., and H.G. designed and conducted experiments. S.D. and M.S.S. provided bioinformatic support and J.B. managed the dyW mouse colony. K.T. synthetized AP-13. F.B, F.V., S.B., and C.R. provided tissues for myoblast isolation that were used in early unpublished experiments contributing to this study. C.F.B., E.L.M., P.S., N.A.D., J.N.F., E.M., and M.A.M. supervised students, interpreted the results and/or edited to the manuscript.

## Materials and Methods

### Mice and animal care

Husbandry and all experimental protocols using mice were performed in accordance with the guidelines established by the animal committee of the Université de Sherbrooke and the Centre Hospitalier Universitaire Sainte-Justine of the Université de Montréal, which are based on the guidelines of the Canadian Council on Animal Care. mdx (Jackson Laboratory, Stock No: 013141), d16 (Jackson Laboratory, Stock No: 024972), and dyW mice (Jackson Laboratory, Stock No: 013786) were generally analyzed at 6-8 weeks of age. Age matched C57BL/6 mice (Charles River, strain 027) were used as wt ctrl animals. Physiological and functional testing of veh or AP-13 treated dyW mice was performed at 5 weeks of age. Female and male mice were included at equal proportions. To ensure optimal access to water and food, cages containing dyW mice were supplied with long-necked water bottles and wet food. For AP-13 or veh treatment, dyW mice were implanted with equilibrated osmotic minipumps (Alzet, model 1004) at the age of 14 days through a small incision at the level of the scapula. Surgery was performed under isoflurane anesthesia. Osmotic minipumps were loaded either with veh (100 μl of 0.9% NaCl) or AP-13 (1mg/kg/day). APJ^EC^KO mice were generated by crossing Cdh5(PAC)CreERT2 (Taconic) and APJ flox mice^38^. To trigger genetic recombination, mice were injected with tamoxifen (Toronto Research Chemicals Inc, T006000-25) dissolved in corn oil (i.p., 2mg) during five consecutive days.

### Muscle regeneration and histology

Muscle injury was induced by injection of 50 µl of 10 µM cardiotoxin (CTX, Latoxan, L8102) from Naja Mossambica into the *tibialis anterior* (TA) muscle of isoflurane-anesthetized animals that were treated with a single dose of buprenorphine for pain management. Following euthanasia by CO2, TA muscles were harvested and embedded in gum tragacanth (Sigma, G1128), snap-frozen in liquid nitrogen chilled isopentane, and stored at -80 °C. 10 µm thick muscle cryosections were stained using hematoxylin and eosin (Sigma, MHS16). Images have been acquired using a Nanozoomer scanner (Hamamatsu, C10730-12). For immunostaining, muscle cryosections were fixed with 4% paraformaldehyde (TCI America, P0018) for 10 min and then permeabilized with 0.5% Triton-X (Sigma, T8787) for 10 min. Sections were blocked in 5% bovine serum albumin (Thermo Fisher Scientific, BP9703100) for at least 1 h at room temperature. Primary antibodies (eMHC, DSHB, F1.652; laminin, Sigma-Aldrich, L9393; dystrophin, DSHB, MANDRA1-7A10; Pax7, DSHB; myogenin, Abcam, ab124800; CD31, Thermo Fisher Scientific, 14-0311-082; fibronectin, Sigma-Aldrich F3648; PDGFRα, R&D systems, AF1062; F4/80, Biorad, MCA497RT; and APJ, ProteinTech 20341-1-AP) were diluted in blocking solution and incubated overnight at 4 °C in a wet chamber. Appropriate secondary antibodies (Thermo Fisher Scientific) and Hoechst (Thermo Fisher Scientific, 62249) were applied for 2 h at room temperature, and mounted using Mowiol (Sigma, 81381) for image acquisition. For Pax7 staining, antigen retrieval using hot 10 mM sodium citrate buffer (Sigma, S4641) supplemented with 0.05% Tween 20 (Bio Basic Canada, TB0560) was performed for 20 min and Fab mouse antigen fragment (Jackson ImmunoResearch, 115-007-003) was added during the blocking step. For visualization, an IgG1 specific secondary antibody was used. For minimal fiber feret analysis, sections were stained with dystrophin and analyzed using the Open-CSAM ImageJ macro^39^.

### Analysis of single-cell RNA-sequencing data

A gene expression matrix of RNA-seq data in FPKM (GEO, GSE143437) from tissue resident endothelial cells was used to generate a gene list with the gene ontology tag GO: 0004930 “G protein-coupled receptor activity” using the Biomart mining tool of Ensembl. From the curated expression matrix containing only G protein-coupled receptor (GPCR) activity-related genes, log2 expression of the top expressed GPCRs with cognate ligands was extracted manually and mapped using GraphPad Prism. Colored Uniform Manifold and Projection (UMAP) plots were generated using Seurat version 4.04 based on single cell sequencing data of TA muscles at 5 dpi^22,40,41^ (GEO: GSE143437). Individual cells were colored based on their expression levels of genes APJ, Pax7 and CD31.

### Apelin-13 Synthesis

Pyr-apelin-13 (AP-13) was synthesized using Fmoc chemistry on solid support as previously described (**Figure S1A**)^24^. Briefly, 2-chloro trityl chloride resin (2-CTC, Matrix Innovation, 2-401-1310) was mixed with a solution of amino acid (Fmoc-L-Phe-OH, 1.2 equiv, Chem-Impex International, 02443), *N,N*-diisopropylethylamine (DIPEA, 2.5 equiv, Chem-Impex International, 00141) in dichloromethane (DCM, Thermo Fisher Scientific, D37-20) overnight at room temperature. After removing excess reagents by filtration, the resin was washed consecutively with DCM, isopropanol (Thermo Fisher Scientific, A416-20), DCM, isopropanol, and DCM for 3 min for each solvent. This washing sequence was used to rinse the resin after every reaction (i.e. capping, deprotection or amino acid coupling). Unreacted groups were capped with a mixture of DCM, Methanol (Thermo Fisher Scientific, A412-20), and DIPEA (3.5:1:0.5) for 1 h. The next amino acid was added to the peptide sequence in two steps: Fmoc deprotection and amino acid coupling. The Fmoc protecting group was removed by treating resin twice with 20% piperidine (Chem Impex International, 02351) in *N,N*-dimethylformamide (DMF, Thermo Fisher Scientific, D119-20) for 10 min. The coupling step was carried out using O-(7-Azabenzotriazol-1-yl)-*N,N,N’,N’*-tetramethyluronium hexafluorophosphate (HATU, 5 equiv, Matrix Innovation, 1-063-0001), amino acid (5 equiv), and DIPEA (5 equiv) in DMF at room temperature for 30 min. These steps were repeated to build the full sequence of Pyr-apelin-13. Cleavage of the peptide from the resin and amino acid sidechain deprotection was carried out using a mixture of trifluoroacetic acid (TFA, Chem Impex International, 00289), triisopropylsilane (TIPS, Oakwook Chemical, S17975), ethanedithiol (EDT, Sigma, 8.00795), and water (92.5:2.5:2.5:2.5). The crude peptide was precipitated in tert-butyl methyl ether (TBME, ACROS Organics, AC378720025). After the supernatant was removed by centrifugation, the peptide was dissolved in 10% acetic acid and the aqueous layer was extracted, filtered, and purified by preparative HPLC (ACE5 C18 column 250 × 21.2 mm, 5 µm spherical particle size). The purity (> 95%) and authenticity of the peptide were confirmed by UPLC-MS Waters (Milford, USA, column Acquity UPLC CSH C18, 2.1 × 50 mm packed with 1.7 μm particles) and high resolution mass spectrometry (electrospray infusion on a maXis ESI-Q-Tof apparatus from Bruker, Billerica, USA).

### Apelin-13 proliferation assay

Cells were maintained in at 37°C in a 5% CO2 incubator. ECs (ATCC, CRL-1730) were cultured in endothelial cell growth medium MV2 (PromoCell, C-22022), MV2 supplement mix (PromoCell, C-39226), and 1% Penicillin-Streptomycin solution (Wisent, 450-201-EL). MuSC-derived myoblast were maintained in growth media containing Ham’s F10 (Wisent, 318-050-CL), 20% FBS (Wisent, 2300040033), 1% Penicillin-Streptomycin solution (Wisent, 450-201-EL), and 2,5 ng/mL bFGF (R&D systems, 3139-FB-025). To assess effects of AP-13 on proliferation, media were replaced by endothelial cell growth medium MV2 (PromoCell, C-22022) without supplement or myoblast growth medium without bFGF. Media were exchanged daily. After 3 days of treatment, cells were fixed with 4% paraformaldehyde (TCI America, P0018) for 10 min and then permeabilized with 0.5% Triton X-100 (Sigma, T8787) for 10 min and stained with Hoechst (Thermo Fisher Scientific, 62249). Images were acquired using a high-throughput Operetta microscope (Perkin Elmer). For each AP-13 concentration, 12 pictures of at least 3 biological replicates were quantified using the Harmony high-content imaging and analysis software (Perkin Elmer).

### Skeletal muscle force measurements

Veh controls and AP-13 treated mice were anaesthetised with an intraperitoneal injection of pentobarbital (30 mg / kg). The *in situ* isometric torque tension was measured on the right hindlimb of anaesthetized mice placed on a 37°C preheated platform of the 1300A whole animal system (Aurora Scientific, Canada). The right hindlimb was first shaved, cleaned with 70% ethanol and fixed above the knee joint with a cone point set screw. Then, the foot was positioned at a 90° angle (neutral position) and stabilized with adhesive onto a footplate attached to a 300C-LR dual-mode lever arm (Aurora Scientific, Canada), allowing the mice to push or pull under stimulation. Once the hindlimb was fixed, two sterile needle electrodes were subcutaneously inserted at either side of the tibial nerve to stimulate the posterior muscles of the lower leg, such as the gastrocnemius and the soleus, which are responsible of ankle plantar flexion. The stimulation current was tuned up to achieve a maximum twitch response, and then the leg was stimulated at different frequencies with a 2-min rest between each stimulation, until reaching the maximum torque tension (mN). For *ex vivo* measurements, EDL muscles were isolated by cutting the proximal and distal tendons, and were placed in an organ bath, maintained at 25°C, and filled with Krebs-Ringer’s solution (137 mM NaCl, 5 mM KCl, 2 mM CaCl2, 24.7 mM NaHCO3, 2 mM MgSO4, 1.75 mM NaH2PO4, and 2 g/l dextrose, pH 7.4) bubbled with carbogen (95% O2, 5% CO2). The proximal tendon was fixed in a stationary clamp with a 3-0 suture (Harvard Apparatus, St. Laurent, Canada), and the distal tendon was connected to a dual-mode level arm system 300C-LR (Aurora Scientific, Inc., Aurora, ON, Canada) that provided control of force and positioning of the motor arm. First, the muscle was initially set at a resting tension of 10 mN for 10 min. Then, the stimulation was delivered by a pair of platinum electrodes located on either side of the muscle using supramaximal 0.2 ms square wave pulses. Muscles were stimulated at different frequencies with 2 min rest between contraction to reach maximum isometric tetanic force (P0). The force generated by the muscle was measured and analyzed with a LabView-based DMC program (Dynamic Muscle Control and Data Acquisition; Aurora Scientific, Inc.). Optimal muscle length (L0) was defined as the muscle length at which the maximal twitch force was elicited. The optimum fibre length (Lf) was determined by multiplying L0 by predetermined Lf/L0 ratios: 0.44 for EDL. The cross-sectional area (CSA) of muscle samples was then determined by dividing muscle mass (mg) by the product of Lf and 1.06 mg / mm3, the density of mammalian muscle. P0 values were normalized to the muscle cross-sectional area.

### Fitness tests

Mice were tested using rotarod (Bioseb, LE8505) at a speed of 15 rounds per min. Grip tests were performed using a horizontal single beam and a grid engineered in-house. In order to calculate the normalized mean impulse, the time until task failure was measured and multiplied by the animals body mass. For each test, three measurements have been recorded and the animals were rested for 5 min between each measurement.

### Apoptosis assay

TA muscles were harvested and snap frozen in liquid nitrogen. Frozen samples were thawed on ice, weighted, and homogenized using a Potter-Elvehjem tissue grinder on ice in ice-cold incomplete radioimmunoprecipitation assay buffer (RIPA) containing 50 mM Tris-HCl pH 7.4 (Sigma-Aldrich, T1503) and 100 mM NaCl (Millipore, SX0420) supplemented with 1 mM 1,10-ortho-phenanthroline (Sigma, P9375), 10 μM 3,4-dicholoroisocoumarin (Sigma, D7910), 10 μM leupeptin (Sigma, L8511), and 10 μM E-64 (Sigma, E3132) for protease inhibition. 20 μl RIPA buffer was used per mg of tissue and the RIPA buffer was completed by addition of 0.1% sodium dodecyl sulfate (Thermo Fisher Scientific, BP166), 0.5% sodium deoxycholate (Sigma, D6750), and 1% Nonidet P-40 (Roche, 11754599001) to obtain a final volume of 30 μl. After lysis on ice for 1 h, cellular extracts were centrifuged at 18,000g for 15 min and the supernatants were recovered. 90 μg of protein was used for western blotting with cleaved caspase-3 (Cell Signaling Technologies, 9661), actin (Sigma, A3853), HRP-conjugated anti-mouse (Cell Signaling Technologies, 7076), and HRP-conjugated anti-rat (Cell Signaling Technologies, 7074) antibodies. Chemiluminescence was acquired with a VersaDoc 4000mp imaging system (BioRad) using the Immobilon Crescendo Western HRP substrate (Millipore, WBLUR0500) or Clarity Max Western ECL Substrate (BioRad, 1705062).

### Echocardiography

Morphological and functional heart parameters have been assessed using a Vevo3100 echocardiography system equipped with a MX400 ultrasound probe (FUJIFILM VisualSonics). Animals were anesthetized using isoflurane and parameters have been recorded at a heart rate near to 450 beats per minute. Cardiac function measurements were acquired in M-mode from a parasternal short-axis view of the left ventricle and analysed using Vevo LAB 3.1.1 (FUJIFILM VisualSonics).

### Quantification and statistical analysis

Except for animals that died a natural death during the course of the experiments, no mice were excluded from the study. Sample size determination was based on the expected effect size and variability that was previously observed for similar readouts in the investigators laboratories. *In vivo* treatments were not blinded, but imaging readouts were analyzed in a blinded manner. Stained samples were analyzed using n≥3 images per biological replicate. Statistical analysis was performed using GraphPad Prism (GraphPad Software). Statistical significance for binary comparisons was assessed by a student’s t-test after verification that variances do not differ between groups or by a Welch correction when variance was observed. For comparison of more than two groups, one-way or two-way ANOVAs were used, according to the experimental design, and followed by Tukey or Dunnett’s post-hoc test. All data are expressed as means + sem.

## Supplementary Information

**Supplementary Fig. 1:**
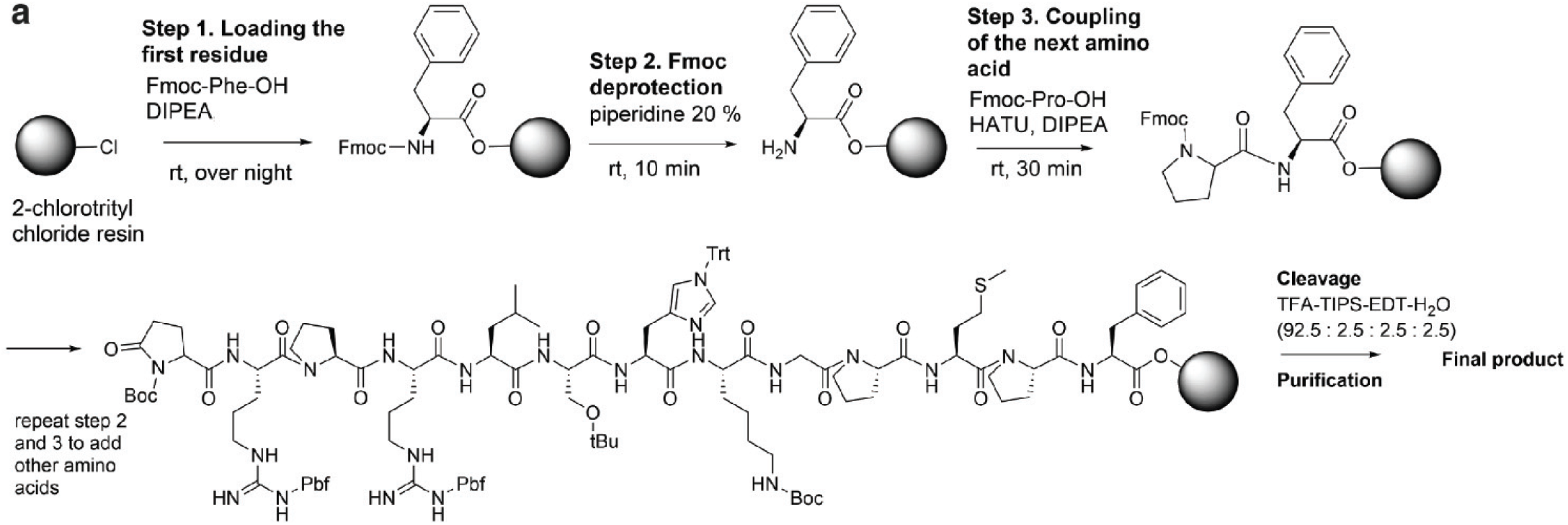
AP-13 synthesis. **a**, Experimental scheme depicting the chemical synthesis of AP-13. rt = room temperature, Fmoc = 9-Fluorenylmethoxycarbonyl, DIPEA = *N,N*-diisopropylethylamine, HATU = O-(7-Azabenzotriazol-1-yl)-*N,N,N’,N’*-tetramethyluronium hexafluorophosphate, TFA = Trifluoroacetic acid, TIPS = triisopropylsilane, EDT = ethanedithiol.

**Supplementary Fig. 2:**
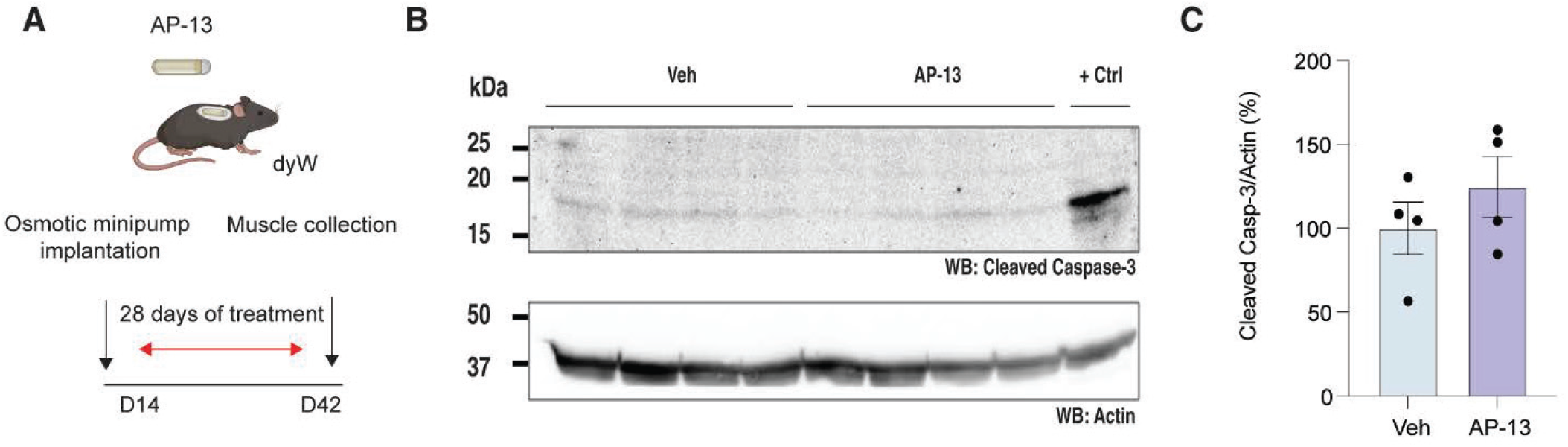
Apoptotic profiling following AP-13 treatment. **a**, Scheme outlining the treatment strategy of dyW mice with vehicle (veh) or AP-13. **b,c**, Western blot (b) and gray value quantification (c) of cleaved caspase-3 (top) and actin (bottom) proteins from skeletal muscle tissues treated with veh or AP-13. +Ctrl = positive control using regenerating C57BL/6N wild-type skeletal muscle tissue at 2 dpi. Bars represent means ± sem. n=4 mice per condition. A student *t*-test was used to determine statistical differences.

**Supplementary Fig. 3:**
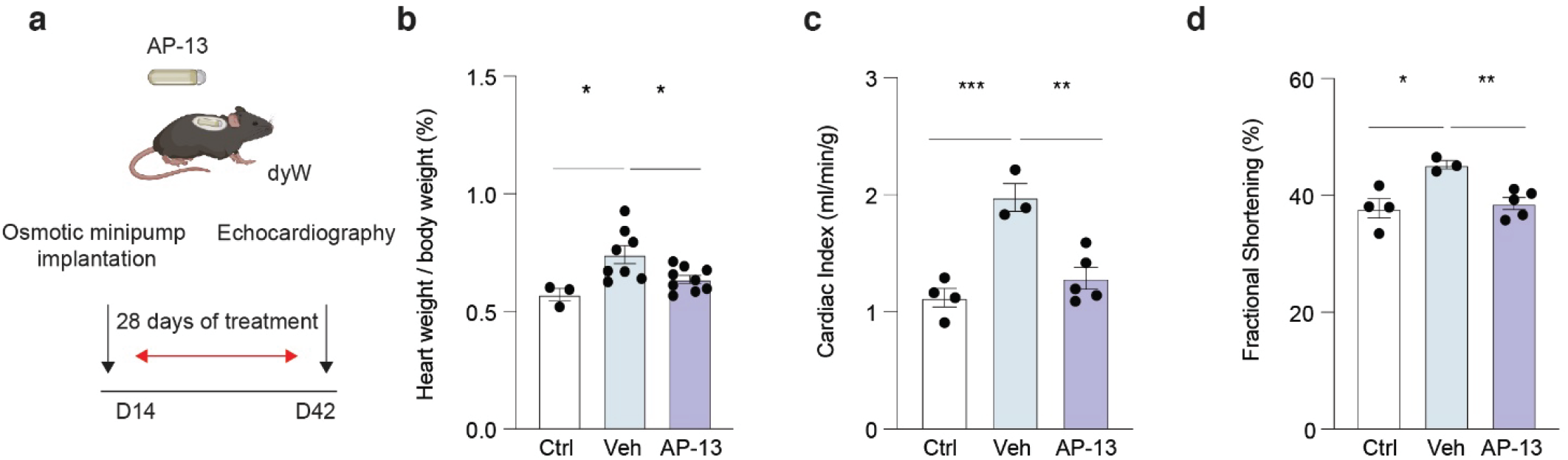
AP-13 does not cause adverse effects on heart function. **a**, Scheme outlining the treatment strategy of dyW mice with veh or AP-13. **b**, Quantification of heart weight normalized to body mass of wt controls (ctrl), veh and AP-13 treated dyW mice. **c**, Quantification of the cardiac index (cardiac output / body mass) of ctrl, and veh and AP-13 treated dyW mice. **d**, Quantification of the heart fractional shortening of ctrl, and veh and AP-13 treated dyW mice. Results are expressed as means ± sem. n≥3 per condition. P values were calculated using one-way ANOVA with Tukey post-hoc test. *P<0.05, **P<0.01, ***P<0.001.

**Supplementary Fig. 4:**
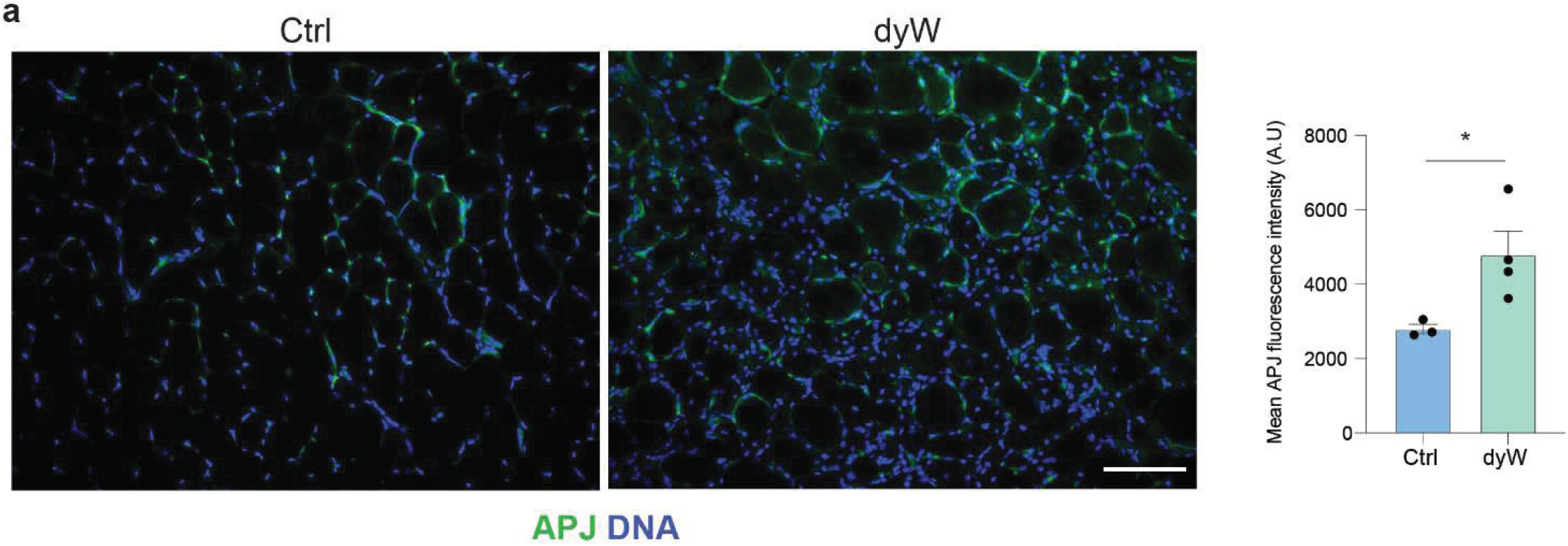
APJ expression in dyW skeletal muscle tissue. **a**, Immunostaining and quantification of APJ expression in TA muscle sections in 6 week old ctrl and dyW mice under uninjured conditions. Results are expressed as means + sem. n≥3 mice per condition. Scale bar = 100 μm. P values were calculated using student *t*-test. \**P*<0.05, \*\**P*<0.01, \*\*\**P*<0.001, \*\*\*\**P*<0.0001.

